# Inhibitory neurosteroid reverses the dendritic spine disorder caused by gain-of-function GABA_A_R epilepsy variants

**DOI:** 10.1101/2021.12.08.471533

**Authors:** Saad Hannan, Kamei Au, Trevor G Smart

## Abstract

GABA_A_ receptors (GABA_A_Rs) are key orchestrators of neuronal activity and several GABA_A_R variants have been linked to genetic neurodevelopmental disorders (NDDs) and epilepsies. Here, we report two variants (Met263Lys, Leu267Ile) in the predominant GABA_A_R α1 subunit gene (*GABRA1)* that increase apparent receptor affinity for GABA and confer spontaneous receptor activity. These gain-of-function features are unusual because GABA_A_R variants are traditionally thought to cause seizures by reducing inhibitory neurotransmission. Both Met263Lys and Leu267Ile increased tonic and spontaneous GABAergic conductances in neurons revealed by competitive inhibition and channel block of GABA_A_Rs. Significantly, α1-subunit variant expression in hippocampal neurons also reduced dendritic spine density. Our results indicate that elevated GABAergic signalling can precipitate genetic epilepsies and NDDs. Furthermore, the mechanistic basis may involve the de-compartmentalisation of excitatory inputs due to the removal of dendritic spines. This aberrant structural plasticity can be reversed by the naturally-occurring, therapeutically-tractable, inhibitory neurosteroid, pregnenolone sulphate.

## Introduction

γ-aminobutyric acid type-A receptors (GABA_A_Rs) mediate inhibitory signalling in the brain. Upon their activation by the brain’s most abundant inhibitory neurotransmitter, GABA, these receptors increase the membrane conductance to Cl^-^ and HCO_3_ ^-^ that collectively causes membrane hyperpolarisation and/or shunting of excitatory synaptic potentials^1,2^. These receptors are known to be vital for controlling neuronal excitability and it is therefore unsurprising that genetic variants of GABA_A_Rs feature prominently in a wide variety of neurological and neuropsychiatric disorders^3,4^. GABA_A_Rs are hetero-pentameric ligand-gated ion channels composed from nineteen subunits (α1-6, β1-3, γ1-3, ρ1-3, δ, θ, ε, π) with prototypical receptors comprised of 2α, 2β, and γ or δ subunits^1,5^.

Genetic epilepsies manifest as sudden uncontrolled bursts of electrical activity in the brain resulting in seizures^6,7^. Subunit variants of nearly all the major isoforms of GABA_A_Rs have been implicated in causing genetic epilepsies that are often co-morbid with neurodevelopmental disorders (NDDs). The mechanisms by which GABA_A_R variants can cause epilepsy ultimately results in dysfunctional inhibition variously achieved by altering: receptor sensitivity to GABA or other ligands^8^; GABA_A_R activation and deactivation kinetics^8–10^; assembly^11,12^; trafficking and cell surface expression^10,13–15^; and degradation^16,17^.

Dysfunctionally low levels of brain neurosteroids are also associated with epilepsy (eg, gender-specific catamenial epilepsy)^18^. This is significant since neurosteroids are potent endogenous modulators of GABA_A_Rs and are tractable compounds for treating GABAergic disorders, including epilepsy^18,19^. Brain neurosteroids can be functionally categorised into two main groups – those that exert positive allosteric and direct activation effects at GABA_A_Rs, such as allopregnanolone, and those that act as negative allosteric modulators, inhibiting GABA_A_R function, such as pregnenolone sulphate (PS).

Although reduced GABAergic signalling is intuitively presumed to initiate seizures and NDDs, it is notable that pathogenic GABA_A_R variants exhibiting increased activity do exist in non-neuronal tissues^20,21^ and are likely to do so in the brain. However, the mechanisms by which increased GABAergic signalling initiates pathogenesis is unknown, and furthermore, there are no therapeutic options for targeting such hyperactive GABA_A_R variants.

The α1-GABA_A_R is the major isoform in the brain accounting for ∼35% of GABA_A_Rs^22^. Two human α1-subunit epilepsy and NDD variants (accession - VCV000280804.2, c.788T>A, p.Met263Lys, Met236Lys; accession - VCV000205521.3, c.799C>A, p.Leu267Ile, Leu240Ile, numbers refer to the mature protein) were selected from the ClinVar database based on their proximity to the positive allosteric neurosteroid binding site located at the receptor’s β-α subunit interface, within the transmembrane domain straddling βM3-αM1 (Fig. 1a-c)^23^.

**Figure 1.**
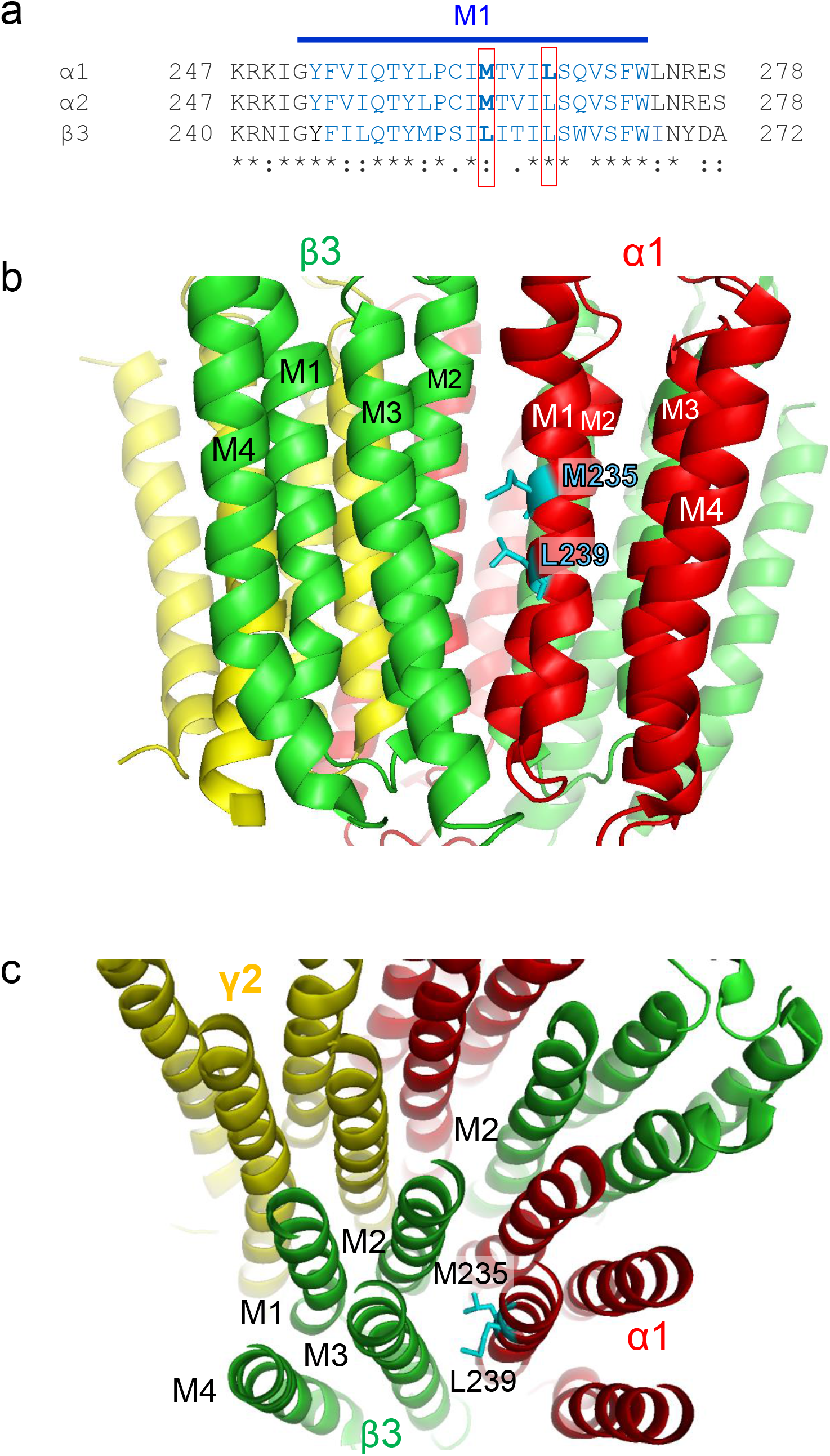
Location of GABA_A_R variants. **a** Primary amino acid sequence alignment of transmembrane domain 1 (M1; residues in blue) for human α1, α2 and β3 subunits. The numbering includes respective signal sequences. The red boxes show two residues: M263 of α1 and α2 (also the equivalent L256 of β3); and L267 of α1. **b, c** 3D structure of an α1β3γ2L GABA_A_R showing the TMD location of α1-M263 and α1-L267 in M1 in side-(**b**) and top-down (**c**) views. The structure was based on PDB 6I53^53^.

Here, we report that these variants unexpectedly confer gain-of-function properties on the receptors with consequences for the structural dendritic plasticity of principle neurons. By examining the profiles of GABA_A_R α1 subunit-containing (α1-GABA_A_R) variants causing epileptic and NDD phenotypes, we probe how a native neurosteroid may be useful in reversing these key detrimental effects.

## Results

### Spontaneously active GABA_A_R epilepsy variants

To explore the biophysical properties of α1 subunits incorporating M236K and L240I, we created recombinant mutant α1 subunits (M235K and L239I) and expressed them in HEK-293 cells as α1β2γ2L assemblies. Generating GABA concentration response curves revealed that the receptors carrying M235K or L239I possessed increased sensitivity to GABA (lower EC_50_s) compared to wild-type counterpart receptors (Fig. 2a-c, P<0.001, One-way ANOVA) with lower maximal currents at saturating GABA concentrations (Fig. 2d; P<0.01/ P<0.001, One-way ANOVA). Another notable feature was the lower Hill slope for M235K. Studying the macroscopic properties of maximal GABA currents for the α1-variants revealed a slower deactivation rate (Fig. 2d; P<0.001, One-way ANOVA) and reduced desensitisation (P<0.01/ P<0.001, One-way ANOVA) without changing receptor activation kinetics (p = 0.3063).

**Figure 2.**
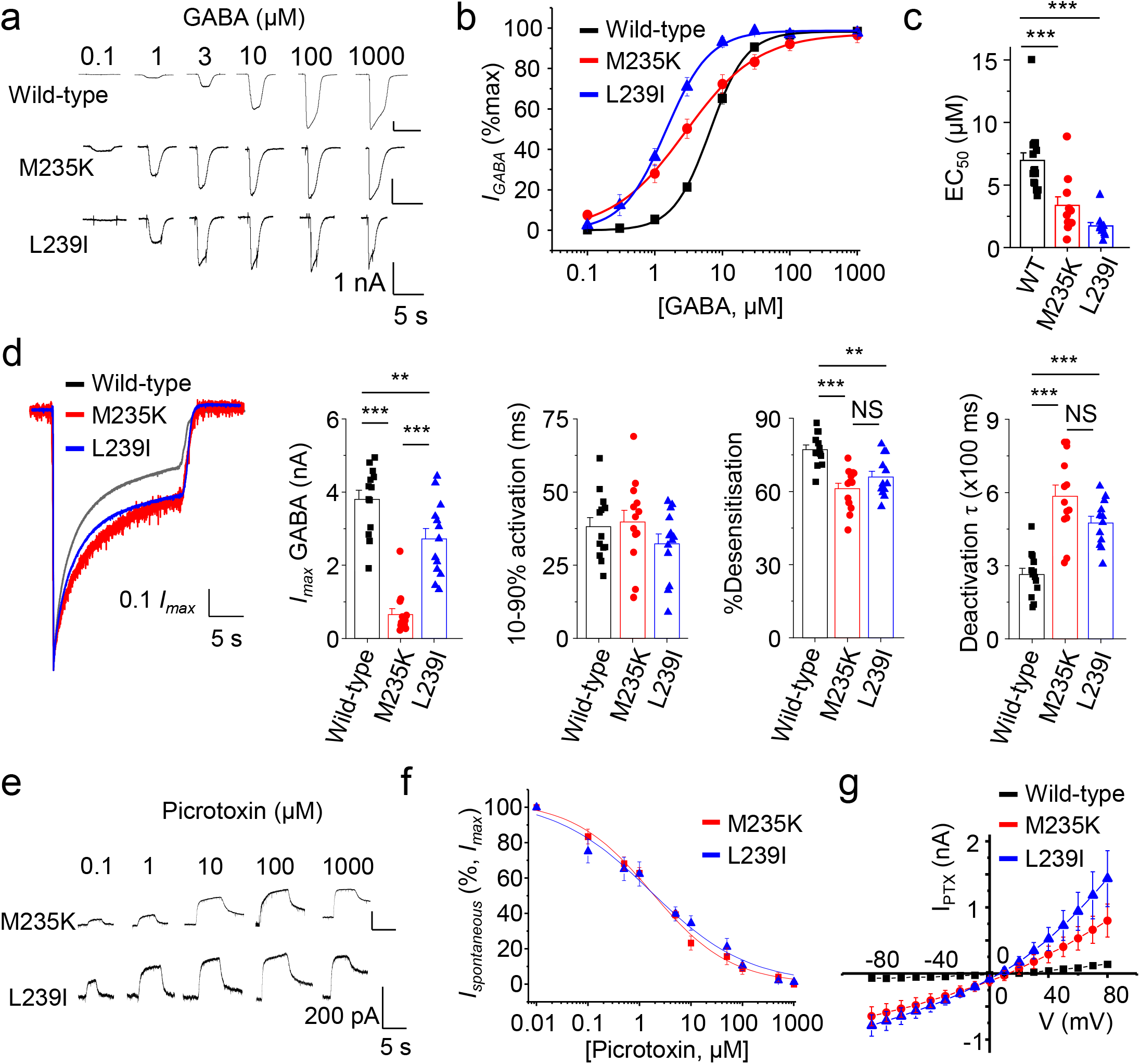
Spontaneous activity of GABA_A_R variants. GABA-activated currents (**a**), GABA concentration response relationships (**b**), and mean GABA EC_50_s (**c**). **d** Macroscopic kinetic properties of GABA currents, left to right panels are: averaged peak-scaled GABA-activated current waveforms evoked by saturating 1 mM GABA, mean 10 – 90 % GABA current activation time, % desensitisation during GABA application and deactivation time constant after GABA washout. **e** Outward currents following concentration-dependent block by picrotoxin of the spontaneous current. **f** picrotoxin inhibition curves for the spontaneous current. **g** Current-voltage (I-V) relationships for picrotoxin-sensitive currents. Data accrued from HEK-293 cells expressing either α1 wild-type or variant α1^M235K^ or α1^L239I^ with β2 and γ2L subunits. In **g**, I-V data represent current subtractions of I-V relationships in the presence and absence of picrotoxin. GABA EC_50_s are: α1^WT^β2γ2L = 6.97 ± 0.61 µM (n = 17), α1^M235K^β2γ2L = 3.38 ± 0.68 µM (n = 11), α1^L239I^β2γ2L = 1.73 ± 0.28 µM (n = 11). Picrotoxin IC_50_s are: α1^M235K^β2γ2L = 1.74 ± 0.37 µM (n = 5), α1^L239I^β2γ2L = 1.67 ± 0.79 µM (n = 5). Bar graphs in this and succeeding figures represent means ± S.E.M. of individual data points (symbols); **P<0.01, ***P<0.001; One-way ANOVA post-hoc Tukey test. *F*_(2, 36)_ = 22.3, p<0.0001 (**c**); *F*_*(2, 36)*_ = 44.3, p<0.0001 (**d**, mean current); *F*_*(2, 36)*_ = 1.22, p=0.3063 (**d**, 10 – 90 % current activation time); *F*_*(2, 35)*_ = 14.6, p<0.0001 (**d**, % desensitisation of peak current); *F*_*(2, 35)*_ = 23, p<0.0001 (**d**, deactivation tau for GABA currents); n = 5 - 17 cells.

The lower maximum currents for M235K and L239I could reflect reduced cell surface expression. However, using antibody labelling in HEK-293 cells, no change was observed for L239I (P>0.05), although cell surface expression of M235K was reduced (Supplementary Fig. 1; P<0.001, One-way ANOVA). The reduced maximum currents and increased potency were not a consequence of aberrant assembly with receptors lacking γ2 subunits^24^, since the expressed α1-variant receptors, and wild-type counterpart, were insensitive to Zn^2+^, which is a ‘fingerprint’ for α1β2γ2 receptors (P>0.05) contrasting with α1β2 receptor currents which are highly-sensitive to Zn^2+^ inhibition (Supplementary Fig. 2; P<0.001, One-way ANOVA)

Under basal GABA-free conditions, cells expressing α1-variants exhibited unusually high leak currents that were reduced by the GABA_A_R channel blocker picrotoxin, in a concentration-dependent manner (Fig. 2e-g). This is indicative of spontaneous activity and was confirmed by current-voltage (I-V) relationships with picrotoxin revealing a basal current at all voltages in the absence of GABA, which remained minimal for wild-type receptors (Fig. 2g). These results indicate that α1^M235K^ and α1^L239I^-GABA_A_Rs are spontaneously-active, more sensitive to GABA, with some limited cell surface expression, and aberrant gating kinetics. Many of these changes are indicative of a gain-of-function profile - this is unusual even counter-intuitive for GABA_A_R epilepsy-inducing variants, which are normally associated with compromised inhibitory signalling.

### Increased tonic currents and spontaneous activity of GABA_A_R epilepsy variants in neurons

To probe the functional consequences of spontaneously-active α1-variants in a native environment, we transfected hippocampal neurons for electrophysiological analysis. Expression of just wild-type α1 subunits did not affect whole-cell muscimol (100 μM) current density (p = 0.2446, selected as a specific GABA_A_R agonist), or the amplitude (p = 0.1519) and frequency (p = 0.3693) of GABA-mediated spontaneous inhibitory postsynaptic currents (sIPSCs), compared to eGFP-expressing or untransfected neurons (Fig. 3a-d; One-way ANOVA). We presume that transfecting the α1 construct did not cause receptor overexpression because the endogenous supply of β and γ subunits is rate limiting. However, sIPSC kinetics in wild-type α1 expressing neurons reduced their decay time (p = 0.0232, p = 0.0007) without changing rise-time (p = 0.131) or charge transfer (p = 0.291) (Fig. 3e; One-way ANOVA) probably because increasing the number of α1 subunits simply substituted for other α-subunit synaptic GABA_A_Rs with slower kinetics. Given that M235K and L239I are spontaneously active, in a native neuronal environment context, these variants should increase the basal inhibitory tone at both synaptic and extrasynaptic locations.

**Figure 3.**
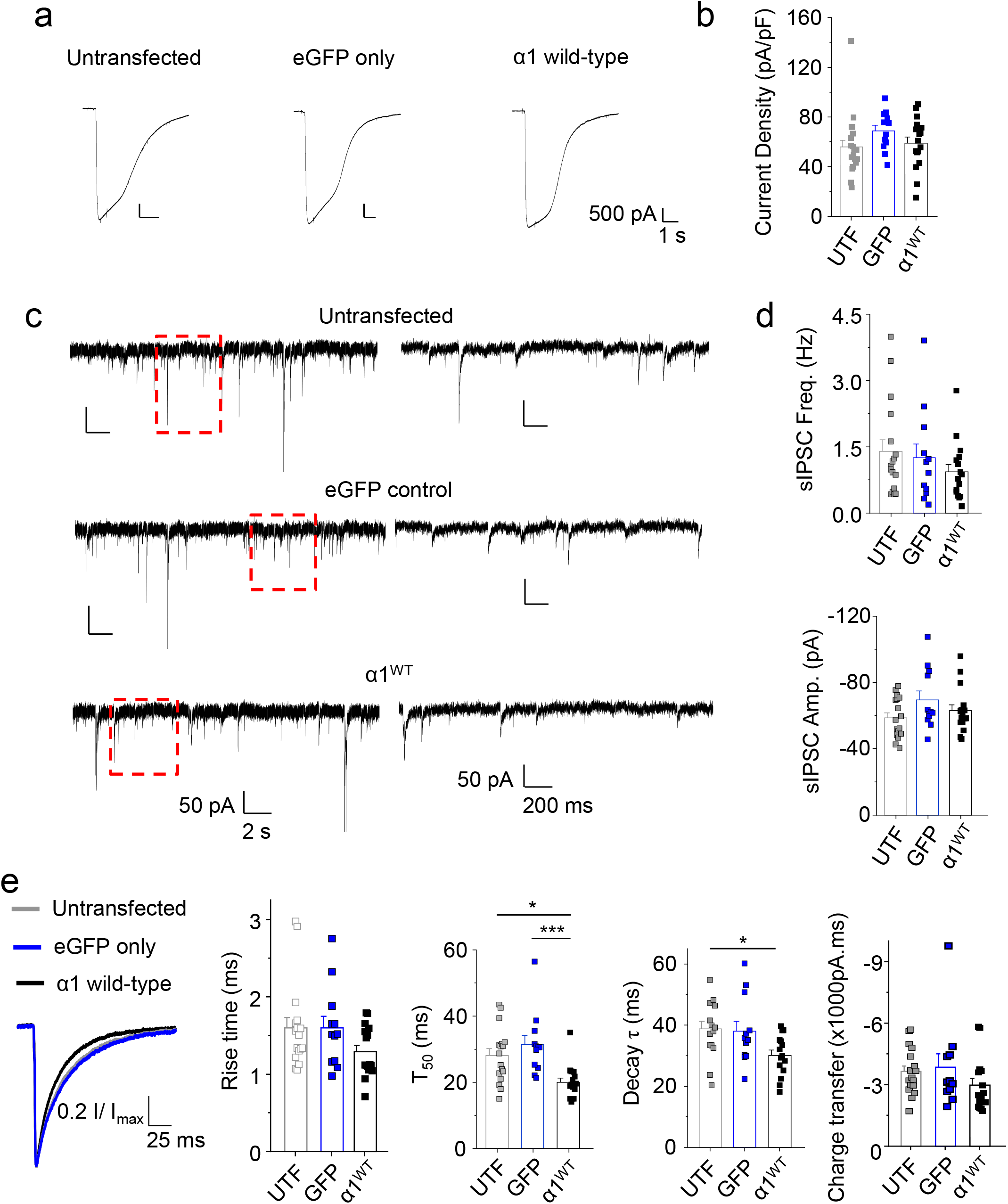
Electrophysiology of α1-GABA_A_Rs expressed in hippocampal neurons. **a** Whole-cell 100 μM muscimol-activated currents recorded at −20 mV from untransfected (UTF), eGFP-expressing neurons, and neurons expressing wild-type α1 and eGFP at 12-14 *DIV*. **b** Mean muscimol current densities of neurons. n = 12 - 21 neurons. *F*_*(2, 47)*_ = 1.45, p = 0.2446. **c** Spontaneous IPSCs recorded from untransfected (UTF) dissociated hippocampal neurons, and from neurons expressing just eGFP (GFP) or with wild-type α1 (α1^WT^) GABA_A_Rs. Higher time resolution records for selected periods (red boxes) are shown on the right. **d** Mean frequency (Freq., upper panel) and amplitude (Amp., lower panel) for sIPSCs. n = 12 - 17 neurons. *F*_*(2, 42)*_ = 1.02, p = 0.3693 for freq, *F*_*(2, 42)*_ = 1.97, p = 0.1519 for Amp. **e** From left to right panels: averaged peak-scaled sIPSC waveforms; sIPSC rise-times (*F*_*(2, 42)*_ = 2.1, p = 0.131); half-decay times (T_50_; *F*_*(2, 42)*_ = 8.6, p = 0.0007); exponential decay times (*F*_*(2, 39)*_ = 4.15, p = 0.0232); and sIPSC areas (charge transfer; *F*_*(2, 39)*_ = 1.3, p = 0.291) of hippocampal neurons expressing eGFP with or without wild-type α1 (α1^WT^) GABA_A_Rs or untransfected neurons. n = 11 - 17 neurons. *P<0.05, ***P<0.001, one-way ANOVA.

Expressing M235K and L239I significantly increased tonic GABA currents by up to ∼10-fold revealed by the GABA_A_R antagonist bicuculline^5^ (Fig. 4a,b; P<0.001, One-way ANOVA), and by co-applying picrotoxin, which resolved an additional tonic component due to the spontaneous activity of M235K (P<0.001) and L239I (P<0.05). This feature was absent (P>0.05; One-way ANOVA) in wild-type α1-subunit expressing neurons. Membrane current variance (noise) for α1^M235K^ and α1^L239I^-expressing neurons was also increased compared to wild-type (P<0.001, One-way ANOVA; Fig. 4c,d), and provided another indicator of increased spontaneous activity. Bicuculline partly reduced the increased variance, which was only normalised to control wild-type levels (p = 0.44, One-way ANOVA) when picrotoxin was subsequently co-applied with bicuculline.

**Figure 4.**
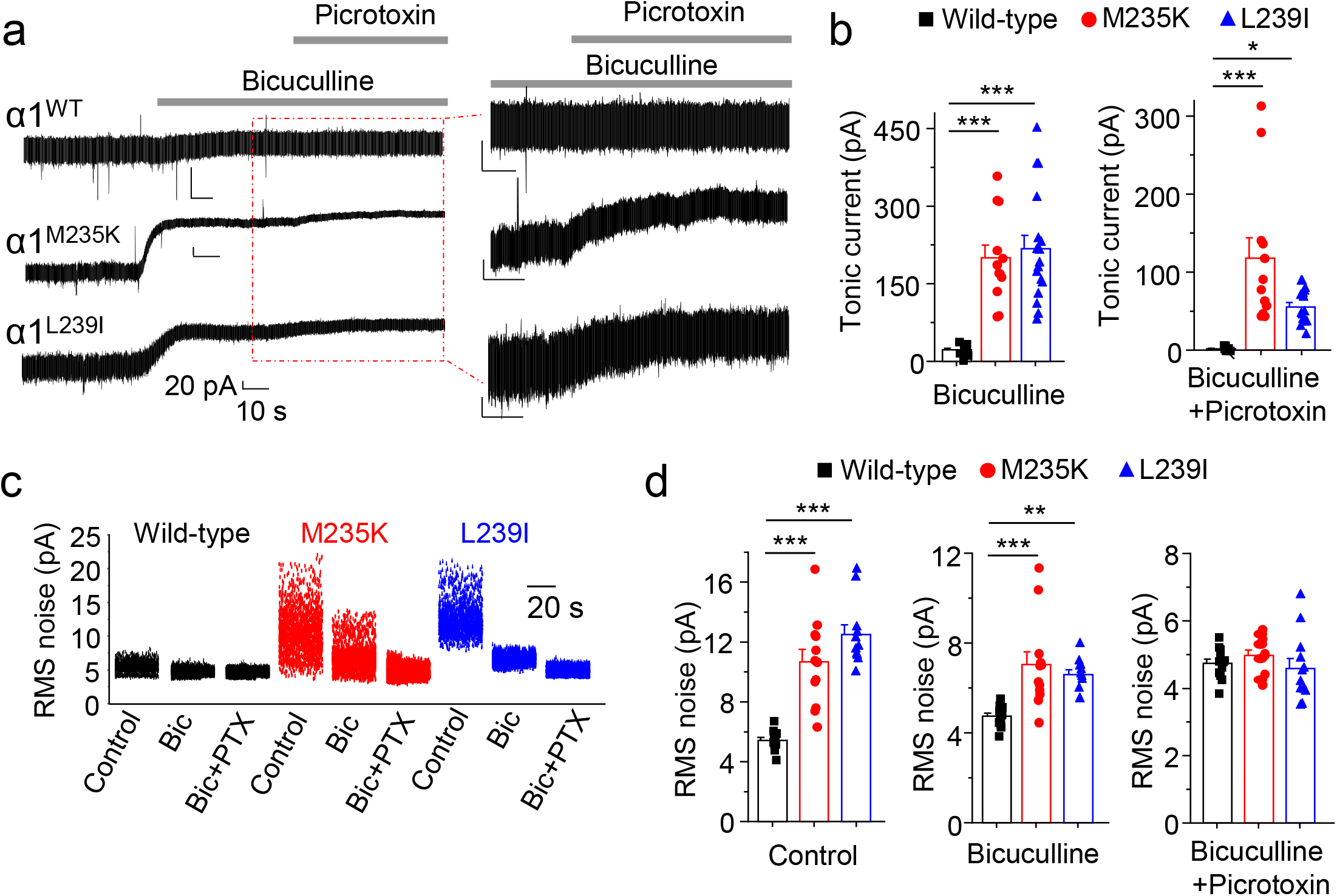
Increased tonic currents and spontaneous activity of GABA_A_R epilepsy variants in neurons. **a** Blockade of tonic and spontaneous GABA-mediated membrane current. Inset (red box) shows the switch between bicuculline (Bic) and Bic + picrotoxin (PTX) at increased resolution. **b** Tonic currents in Bic and in Bic + PTX. **c** Epochs (30 s) of root mean square (RMS) membrane current noise. **d** Comparison of RMS noise for hippocampal neurons expressing wild-type or variant α1 subunits in control or in 25 μM Bic with or without and 100 μM PTX. *F*_(2, 37)_ =19.6, p<0.0001 (**b**, Bic); *F*_(2, 36)_ =14.4, p<0.0001 (**b**, Bic + PTX); *F*_(2, 36)_ = 36.5, p<0.0001 (**d**, control); *F*_(2, 36)_ = 10.5, p=0.0002 (**d**, Bic); *F*_(2, 36)_ =0.84, p=0.44 (**d**, Bic + PTX). n = 11 – 17 neurons. *P<0.05, ***P<0.001, one-way ANOVA.

To ascertain the cell surface expression levels for the α1-variants in hippocampal neurons, N-terminal myc-tagged subunits were used in conjunction with immunocytochemistry. Both α1-variants showed reduced cell surface expression compared to their wild-type equivalents (Supplementary Fig. 3; P<0.01, P<0.001, One-way ANOVA).

Overall, these results suggest that the spontaneously-active α1-GABA_A_R variants are expressed on neuronal surface membranes, albeit at a reduced level, resulting in increased GABA-mediated tonic current, and notably, a spontaneous GABA_A_R-mediated membrane conductance.

### Reduced spine density due to GABA_A_R epilepsy variants

To assess the impact of elevated GABA-dependent and -independent tonic membrane conductances on excitatory synaptic inputs, we measured miniature excitatory postsynaptic currents (mEPSCs) in hippocampal neurons treated with tetrodotoxin. Applying bicuculline and picrotoxin revealed no change to mEPSC frequency (p = 0.94) or amplitude (p = 0.3446) (Fig. 5a,b; One-way ANOVA) implying that, functionally, postsynaptic inputs were apparently unperturbed by either α1^M235K^ or α1^L239I^. However, quite unexpectedly, the structural plasticity of dendritic spines was affected, potentially underlying seizure activity^25^, with spine density reduced (p = 0.0206) without changing the mean spine head diameter (p = 0.6217) for M235K and L239I compared to wild-type GABA_A_Rs (Supplementary Fig. 4d, Fig. 5c,d; One-way ANOVA). Categorising the dendritic spines revealed reduced numbers of mushroom spines (p = 0.0263) and increased thin spines (p = 0.0237), with stubby spines remaining unchanged (p = 0.9816), for neurons expressing the α1-variants (One-way ANOVA; Supplementary Fig 4e, Fig. 5d). These changes to spine density, spine type and size were not apparent on expressing wild-type α1 subunits or eGFP alone, indicating α subunit expression *per se*, and /or subunit-switching artefacts, do not account for the results with the α1-variants (Supplementary Data Fig. 4a-c; P>0.05, two-tailed unpaired t-test).

**Figure 5.**
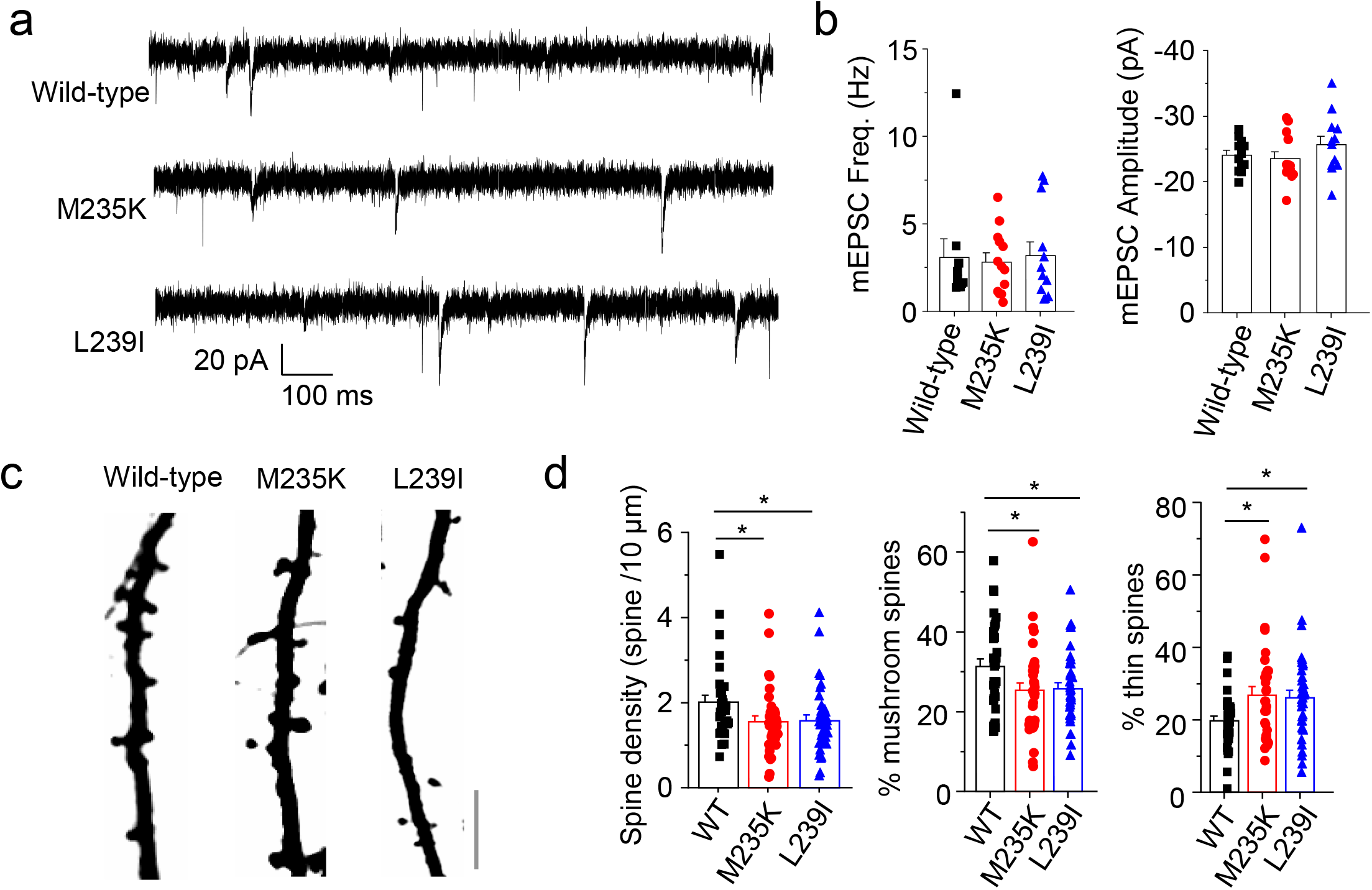
GABA_A_R epilepsy variants reduce spine density without affecting mEPSCs. **a** Miniature EPSCs recorded from dissociated hippocampal neurons expressing wild-type or α1-variant GABA_A_Rs and eGFP at 12-16 *DIV*. mEPSCs were recorded at −70 mV in the presence of 0.5 μM tetrodotoxin, 25 μM Bic and 50 μM PTX. (**b**), Mean frequency (Freq.) and amplitude of mEPSCs expressing wild-type or α1-variant GABA_A_Rs. (*F*_*(2, 32)*_ = 0.06, p=0.94) for frequency and (*F*_*(2, 34)*_ = 1.1, p = 0.3446) for amplitude. **c** Images of dendrites from neurons expressing wild-type or variant α1-GABA_A_Rs with eGFP. **d** Mean spine density and relative proportions (%) of mushroom-shaped and thin spines for neurons expressing wild-type or variant α1-GABA_A_Rs. n = 16 - 36 neurons. *P<0.05, One-way ANOVA. Calibration bars = 5 μm. p = 0.0206 (density); p = 0.0263 (% mushroom); p = 0.0237 (% thin).

### Inhibitory neurosteroids reverse dendritic spine defects due to GABA_A_R epilepsy variants

To attempt to correct the deficits in spine density, we presumed that controlling the increased activity of the α1-variants would be critical. To achieve this, we selected a naturally-occurring inhibitory neurosteroid in the brain, PS, to act as a negative allosteric modulator^26^. Applying 5 µM PS for 48 hr to neuronal cultures expressing α1^L239I^ reversed the deficits in spine density, contrasting with the lack of effect of PS on dendritic spines of neurons expressing just eGFP or wild-type α1 subunits (Fig. 6a-c; p = 0.0136, p = 0.7372, two-tailed unpaired t-test; Supplementary Fig. 4f). For M235K, spine density showed a tendency to increase in PS (p = 0.09) compared to untreated M253K-expressing cells (Fig. 6b). Furthermore, the differences in mushroom, stubby and thin spines between α1-variants and α1 wild-type expressing neurons were absent in the presence of PS suggesting that the neurosteroid affected spine maturation for spontaneously-active GABA_A_R variant-expressing neurons (Supplementary Fig. 5a, P>0.05, One-way ANOVA). Application of PS did not change the frequency or amplitudes of mEPSCs for α1-variant and wild-type α1-subunit expressing neurons (Fig. 6d-f, P>0.05, two-tailed unpaired t-test).

**Figure 6.**
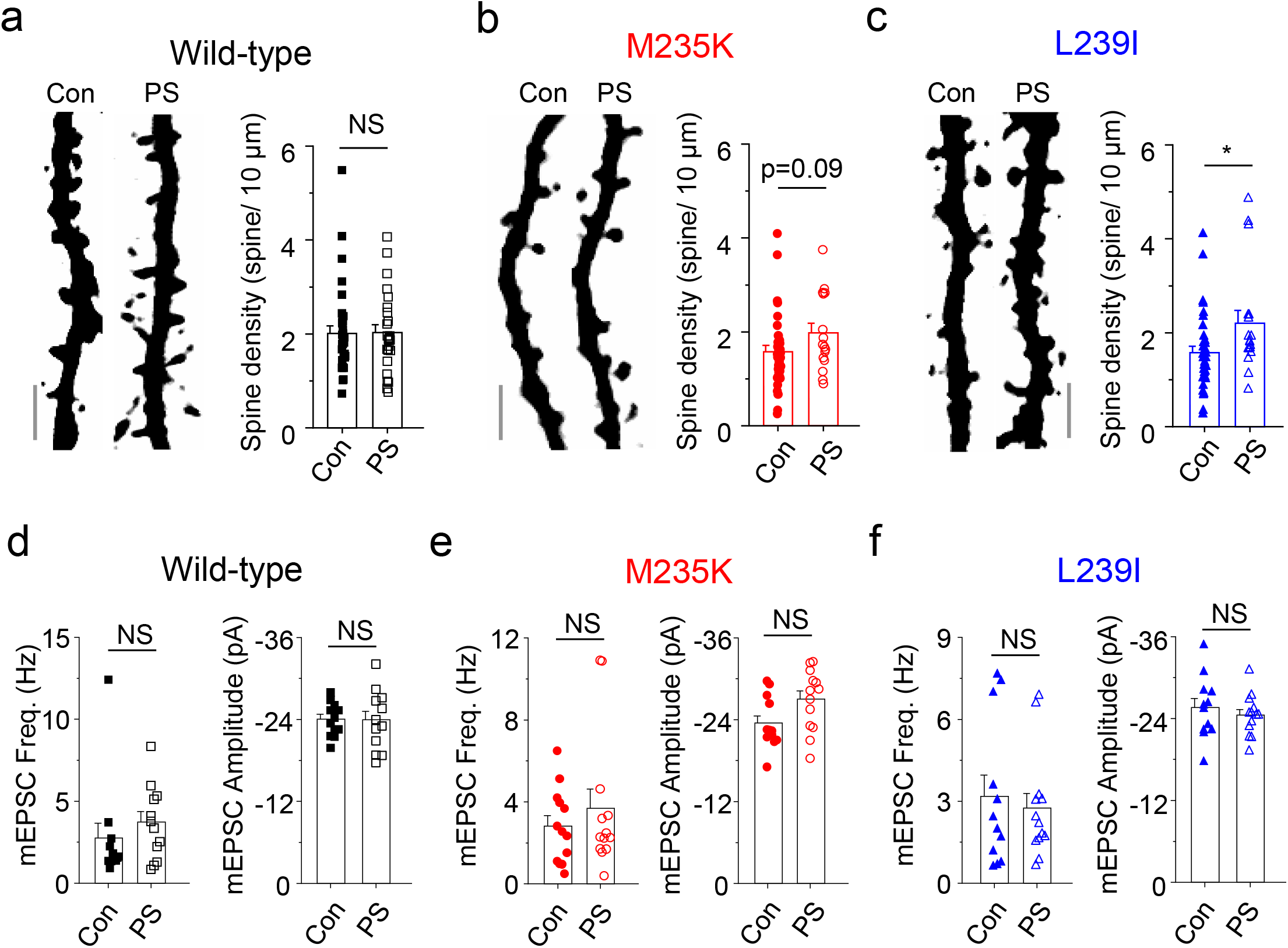
Pregnenolone sulphate reverses spine density defects. **a** Dendritic images and spine density for neurons expressing wild-type α1-GABA_A_Rs and eGFP in control or after 48 hr at 37°C of 5 μM pregnenolone sulphate (PS). **b** Dendritic images and spine density of neurons expressing α1^M235K^-GABA_A_Rs and eGFP in control or after PS treatment as in **a. c** Dendritic images and spine density of neurons expressing α1^L239I^-GABA_A_Rs and eGFP in control or after 48 hr in PS. n = 16 - 36 neurons. NS – not significant, *P<0.05, two-tailed unpaired t test, Mann-Whitney test, Calibration bars = 5 μm. **d** Mean mEPSC frequency (p = 0.4025) and amplitude (p = 0.9355) for wild-type α1-expressing neurons in control and after PS. Neurons were treated with 5 μM PS for 48 hr at 37°C prior to imaging. **e** Mean mEPSC frequency (p = 0.421) and amplitude (p = 0.0538) for α1^M235K^-expressing neurons in control and in PS. **f** Mean mEPSC frequency (p = 0.9362) and amplitude (p = 0.5033) for α1^L239I^-expressing neurons in control and in PS. n = 12 - 13 neurons. NS – not significant; two-tailed unpaired t-test.

The significance of reducing spontaneous GABA_A_R activation for correcting spine deficits was evident with the more potent GABA antagonist, picrotoxin (50 µM). When applied for the same duration as PS, picrotoxin increased spine density for α1^M235K^ (p = 0.014) and α1^L239I^ (p = 0.0007), as well as for eGFP only (p = 0.0082) expressing neurons (Fig 7a,c,d; two-tailed unpaired t-test). There was also a trend for spine density of wild-type α1-expressing neurons to increase in picrotoxin (Fig. 7b) and a similar normalisation of mushroom and thin spines in picrotoxin was evident (Supplementary Fig. 5b; P>0.05, One-way ANOVA) highlighting a common thread of reduced inhibition favouring spine development and/or maintenance. Spine deficits and their reversal by picrotoxin has also been reported with a trafficking defective β3-GABA_A_R subunit^27^.

**Figure 7.**
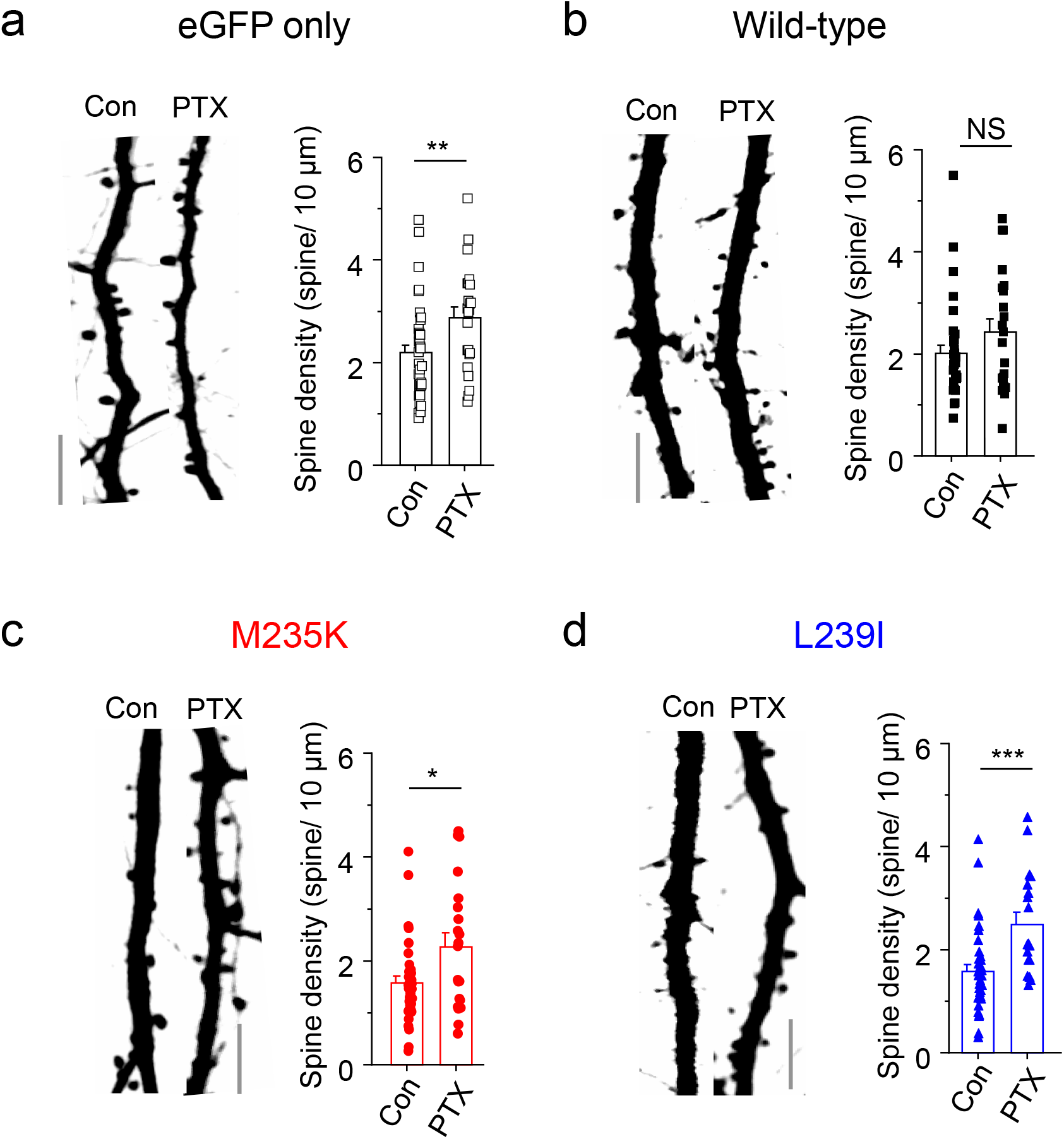
Picrotoxin increases spine density in hippocampal neurons. **a** Confocal images of dendrites from hippocampal neurons expressing just eGFP in control (con) and after 50 µM picrotoxin (PTX) for 48 hr at 37°C. Bargraph shows mean spine density in control and in PTX (p = 0.0082). **b** Confocal images of dendrites from hippocampal neurons expressing α1 wild-type and eGFP in control and in PTX. Bargraph presents mean spine densities (Con vs +PTX, p = 0.1461). **c** Confocal images of dendrites from hippocampal neurons expressing α1^M235K^ and eGFP (Con and + PTX). Mean spine density is increased by PTX (p = 0.014). **d** Images of hippocampal neuronal dendrites expressing α1^L239I^ and eGFP (con and + PTX). Mean spine density is increased by PTX (p = 0.0007). n = 18 - 40 neurons, NS – not significant, *P<0.05, **P<0.01, ***P<0.001, two-tailed unpaired t test. Calibration bars = 5 μm.

### An M1 hotspot for spontaneous active epilepsy-inducing GABA_A_R variants

Finally, we studied two additional variants at the M1 TMD M235 site that are linked to West syndrome severe epilepsy and intellectual disability (ID) ^28,29^. These variants (human M263I (accession - VCV000402327.1) and M263T, numbered by including the signal sequence and equivalent to mouse M235I and M235T in the mature protein) also showed gain-of-function properties increasing GABA potency when expressed as α1β2γ2L receptors, by ∼4 to 14-fold compared to wild-type (Fig 8a-c). In addition, these receptors exhibited reduced maximal GABA currents, and spontaneity, revealed by picrotoxin, suggesting that the TMD methionine residue is crucial for GABA_A_R function (Fig. 8d-e; P<0.001, One-way ANOVA). This region of M1 could therefore represent a critical region (hotspot) for disease-relevant spontaneously-active GABA_A_Rs.

**Figure 8.**
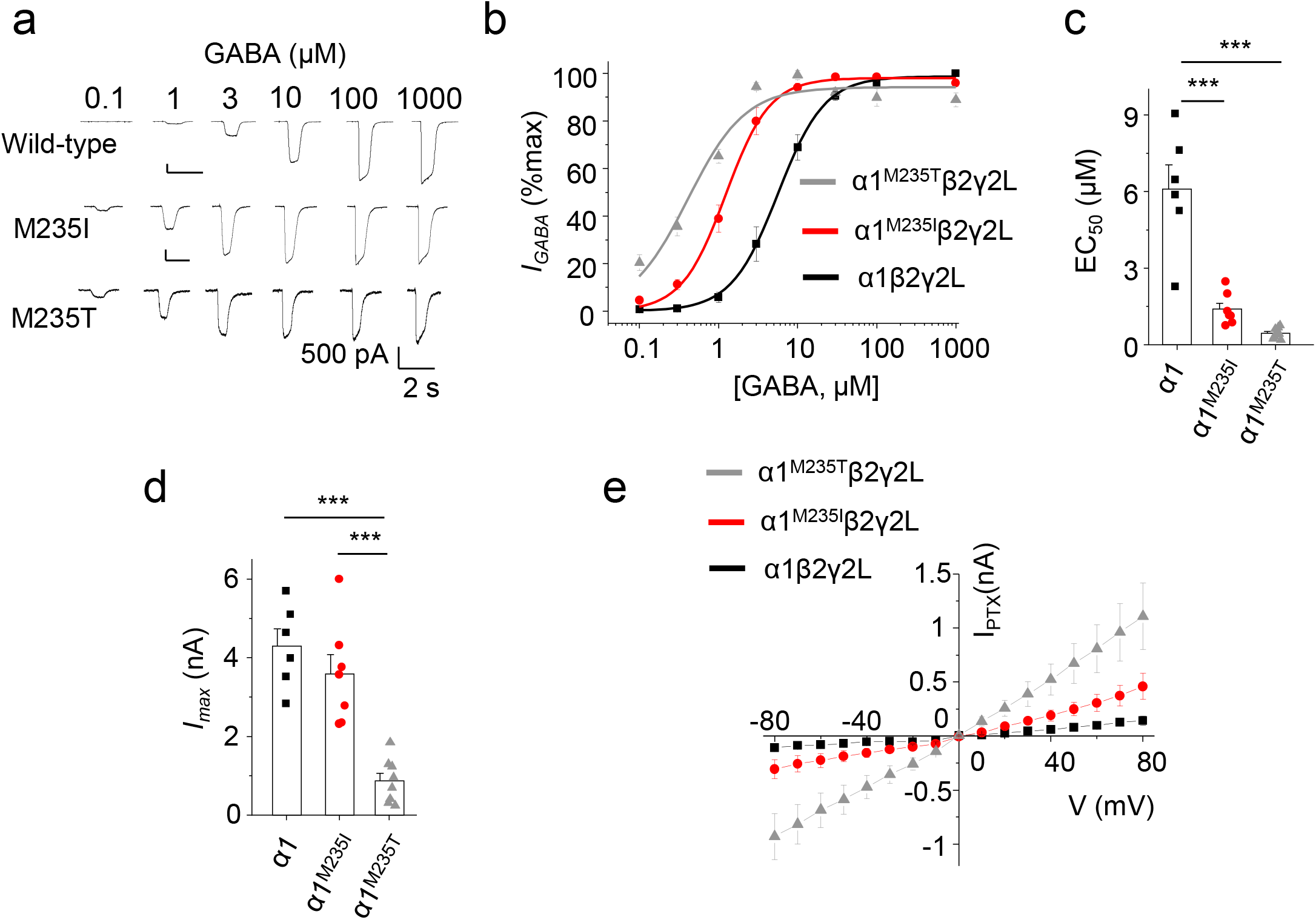
Spontaneous activity of α1^M235I^ and α1^M235T^ variant GABA_A_Rs. **a** GABA-activated currents recorded from HEK-293 cells expressing wild-type α1, α1^M235I^ or α1^M235T^ with β2 and γ2L. **b** GABA concentration response relationships for α1β2γ2L, α1^M235I^β2γ2L and α1^M235T^β2γ2L receptors. **c** Mean GABA EC_50_s for α1β2γ2L, α1^M235I^β2γ2L and α1^M235T^β2γ2L. EC_50_s: α1^WT^β2γ2L = 6.1 ± 0.9 µM (n = 6), α1^M235I^β2γ2L = 1.4 ± 0.2 µM (n = 7), α1^M235T^β2γ2L = 0.45 ± 0.06 µM (n = 9). *F*_*(2, 19)*_ = 40, p<0.0001. **d** Maximal GABA currents for α1β2γ2L, α1^M235I^β2γ2L and α1^M235T^β2γ2L. *F*_*(2, 18)*_ = 23.8, p<0.0001. **e** *C*urrent-voltage (I-V) relationships for PTX-sensitive currents recorded from α1β2γ2L, α1^M235I^β2γ2L and α1^M235T^β2γ2L. I-V curves presented are subtractions of I-V relationships in the presence and absence of PTX. n = 6 - 9, ***P<0.001, One-way ANOVA.

## Discussion

Here, we identify the first α1-GABA_A_R variants to exhibit spontaneous receptor activity that are linked to severe neurological consequences. These spontaneously-active GABA_A_R epilepsy variants are expressed on neuronal plasma membranes and increase GABA-mediated tonic and spontaneous membrane Cl^-^ conductances. The increased tonic GABA current mediated by the α1-variants is likely to reflect the higher apparent affinity these receptors have for GABA, thus increasing their activation compared to wild-type receptors, at ambient extrasynaptic GABA concentrations.

Our results tentatively identify a variant hotspot at the base of α1-GABA_A_R in M1 that generates spontaneously active GABA_A_Rs. The presence of individuals with epilepsy due to substitution of M235 (M235I, M235T and M25K) and neighbouring L239 close to the positive allosteric neurosteroid binding site makes this previously well-characterised area^23,30^ a focus of pathological interest. Notably, equivalent variants located on human GABA_A_R α2 (accession–VCV000689389.2, M263T) and β3 subunits (accession - VCV000975911.1; L256Q) are also linked to severe epilepsy with ID^31,32^. The M1 domains of α1, α2 and β3 subunits are highly conserved (Fig 1a) and α2- and β3-GABA_A_Rs are also major GABA_A_R isoforms in the cortex^22^. Although the functional properties of β3^L256Q^ are unknown, α2^M263T^ increases GABA potency, copying the profile of the α1-variants studied here, suggesting that these α2-receptors may well also increase tonic inhibition. Therefore, the base of the α-helical M1 could represent a critical receptor sub-domain for pathogenic variants initiating neurodevelopmental disorder via spontaneously-active gain-of-function GABA_A_Rs. Potentiating neurosteroids are known to bind to this area^23,30^ and allosterically modulate GABA_A_Rs by increasing GABA potency and facilitating receptor gating^33,34^. Substitution of the identified methionine or leucine residues in this area could alter the packing of the transmembrane domain to enable spontaneous GABA-independent gating. It is also likely that other disparate transmembrane domain variants of GABA_A_R subunits could also achieve similar results by destabilising the ion channel activation and desensitisation gates.

Whereas reduced GABAergic signalling features prominently in the causation of epilepsy and NDD, counterintuitively, our results show that increased apparent affinity and spontaneous activity of GABA_A_Rs can be pro-convulsive therefore sub-classifying this type of hyperactive-GABAergic-dependent epilepsy. The mechanism(s) by which spontaneous activity causes convulsions could involve several possibilities: increased spontaneity could raise intracellular Cl^-^ shifting the equilibrium potential for GABA to depolarising levels or increased membrane shunting of interneurons could reduce GABA release to dampen the inhibition of excitatory networks. Whilst these remain as possibilities, a loss of dendritic spines due to spontaneous GABA_A_R activity as observed in our study, altering neural connectivity to favour excitation over inhibition was not expected. The removal of dendritic spines may result in an inability of neurons to compartmentalise their excitatory inputs. Losing dendritic spines, but without changing mEPSC frequency, suggests that overall neural connectivity remains intact. However, a higher proportion of inputs to α1-variant expressing neurons will likely be made to regions of dendrites that now lack physical compartmentalisation normally afforded by mature dendritic spine structure^35^. Although the precise mechanism remains unknown, a reduced electrical compartmentalisation exacerbated by GABA_A_R activity, could increase electrotonic communication amongst juxtaposed excitatory synapses perhaps leading to the generation of backpropagating action potentials and increased excitability.

Dendritic spines receive the bulk of excitatory inputs and structural plasticity of spines has been studied in detail^36^. For instance, stabilisation of spines has been described to be important in memory and learning^37^. Consistent with this, long-term neural plasticity changes have been reported with picrotoxin treatment in a Down syndrome mouse model of cognitive disability. Picrotoxin was applied to combat excessive GABAergic inhibition in this model^38^ and was accompanied by a reduced dendritic spine density^39,40^.

Interestingly, elevation of tonic GABA-mediated inhibition plays an important role in setting the window for critical period plasticity^41^ and brain circuit development, and here too, dendritic spines undergo dramatic structural changes during critical phases^42,43^ coinciding with the time-point for the maturation of parvalbumin interneurons^44,45^. Furthermore, activation of GABA_A_Rs with agonists such as muscimol, or uncaging of GABA to brain slices, also reduces dendritic spine motility^46^.

The present study has interesting parallels with multiple previous studies that have noted dendritic spine density changes evident in epileptic brain tissue specimens from human and animal models^25,47,48^. In patients with temporal lobe epilepsy, a reduction of dendritic spine density of hippocampal principal neurons has been widely reported. In addition, animal models of chronic and acute seizures also show similar reductions of dendritic spines often followed by formation of varicose swellings^25^. Increasing intracellular Cl^-^ levels have been held responsible for the formation of varicose bodies during excitotoxic insults^49,50^. The intriguing question is what is the role of the spines and why is their removal precipitating epilepsy? By virtue of their high spine neck resistance and low capacitance^51^, spines are considered to normalise the variability of excitatory transmission providing consistency to EPSPs including spike initiation in the dendrite arbour^52^. On this basis, we would expect EPSPs emanating from spine synapses to be faster and of shorter duration compared to dendritic shaft synapses where broader EPSPs would be expected with potential ramifications for integrating excitatory transmission over the dendritic arbour. Our results suggest the loss of spines will exacerbate this difference in excitatory transmission and may underlie the impact that elevated GABAergic signalling has on initiating seizures.

Our results also provide proof-of-concept for using inhibitory neurosteroids to reverse the structural dendritic deficits caused by the mechanistically “atypical” hyperactive GABAergic-dependent epilepsy. Application of a more potent antagonist, picrotoxin, also reversed the spine deficits confirming the role of α1-variant gain-of-function receptors in this form of epilepsy. Interestingly, picrotoxin increased spine density of eGFP-only expressing neurons, as well as for the α1-variants, whereas PS selectively restored dendritic spine levels just for the α1-variant expressing neurons. The spine density increase was greater for picrotoxin compared to PS possibly reflecting the differential potencies for inhibiting GABA_A_Rs. Overall, PS, or alternative inhibitory neurosteroid derivatives, may offer a highly attractive therapeutically-tractable drug alternative for treating such gain-of-function GABA_A_Rs that are associated with epilepsy and NDD.

## Data Availability

The data that are presented in this study are available from the corresponding author on reasonable request. A source data file containing raw data is also included and uploaded online. See further details in the Reporting Summary.

## Acknowledgements

This work was supported by the MRC and Wellcome Trust (TGS), and by an early career fellowship (SH) from the International Rett Syndrome Foundation.

## Contributions

SH conceptualised, and SH and TGS designed and planned the study, SH and KA performed the data acquisition and analysis, TGS performed the receptor modelling, SH and TGS secured project funding. All the authors participated in the writing, reviewing and editing of the manuscript.

## Ethics Declarations

The authors declare no competing interests or conflicts of interest

## Online Methods

### Hippocampal neurons and cell culture

All animal-based studies were performed in accordance with the UK Animals (Scientific Procedures) Act 1986. Cell culture reagents are from ThermoFisher, unless stated otherwise. Embryonic day 18 (E18) Sprague-Dawley rat hippocampi of either sex were prepared and seeded onto 18-22 mm glass coverslips (VWR) coated with poly-D-lysine in minimum essential media with 5% v/v fetal calf serum (FCS), 5% v/v horse serum, 50 units/ 50 µg/ml penicillin-G/ streptomycin, 2 mM glutamine and 20 mM glucose. After 3 hr, the medium was replaced with Neurobasal-A supplemented with 1% v/v B-27, 25 units/ 25 µg/ml penicillin-G/streptomycin, 0.5% v/v Glutamax and 35 mM glucose. Neurons were transfected 6-7 days *in vitro* (DIV) using either a calcium phosphate^54^ or Effectene-based (Qiagen) method.

HEK-293 cells were grown at 37°C in 95% air/5% CO_2_ in Dulbecco’s modified Eagle’s medium supplemented with 10% v/v FCS, 50 units/ ml penicillin-G, 50 μg/ ml streptomycin and 2 mM glutamine. HEK-293 cells were plated onto 22 mm glass coverslips, coated with poly-L-lysine, and transfected 1-2 hr after plating using a calcium phosphate method^55^ with equimolar ratios of cDNAs encoding GABA_A_R α1, β2, γ2L and eGFP subunits.

### cDNA and constructs

cDNAs for wild-type mouse α1, β2, γ2L, α1^myc^ and eGFP have been described previously^11^. Human M263K (with signal sequence) or M236K (mature protein) and murine M235K (equivalent without signal sequence) were created using a single inverse PCR^56^ and ligation using AGACAGTTATTCTCTCCCAAGTCTCC (forward primer) and TTATGCACGGCAGATATGTTTGAATAAC (reverse primer). Human M263I (with signal sequence) or M236I (mature protein) and murine M235I (equivalent without signal sequence), and human M263T (with signal sequence), or M236T (mature protein, and murine M235T (equivalent without the signal sequence) were created with TCACAGTTATTCTCTCCCAAGTCTCCTTC and CGACAGTTATTCTCTCCCAAGTCTCCT as forward primers, respectively, using the same reverse primer as for M235K. For L267I, L240I and L239I, these were created using the same strategy with ATCTCCCAAGTCTCCTTCTGGCTCAACAG (forward primer) and AATAACTGTCATTATGCACGGCAG (reverse primer). The integrity of all cDNAs was confirmed by DNA sequencing.

### Immunolabeling and confocal microscopy

Cells were washed in phosphate-buffered saline (PBS) before fixation in 4% paraformaldehyde for 10 min at room temperature. Myc-tagged α1-GABA_A_R were labelled with mouse anti-myc antibody (ab32; Abcam) followed by goat anti-mouse Alexa Fluor-555 (A28180; ThermoFisher). Cells were imaged at 8-bit immediately following immunolabeling using a Zeiss LSM 510 microscope with a x40 objective and a 488 nm laser for imaging eGFP and a 543 nm laser for imaging Alexa Fluor 555 at optimum z-stack thickness. Images were analysed using Image J (ver 1.52i) by measuring mean cell surface fluorescence levels of defined regions-of-interest drawn around the periphery of cells^54^.

### Electrophysiology

Whole-cell electrophysiology was carried out using borosilicate thin-walled glass patch electrodes (resistances of 3 – 5 MΩ) with optimised series resistance (Rs, <10 MΩ) and whole-cell membrane capacitance compensation. Membrane currents were filtered at 5 kHz (−3 dB, 6th pole Bessel, 36 dB per octave). Cells were superfused with a saline solution containing (in mM): 140 NaCl, 4.7 KCl, 1.2 MgCl_2_, 2.52 CaCl_2_, 11 Glucose, and 5 HEPES; pH 7.4. HEK-293 cells were studied 48 hr after transfection by voltage clamping at −20 to −30 mV using an internal solution containing (mM): 120 CsCl, 1 MgCl_2_, 1 CaCl_2_, 11 EGTA, 30 KOH, 10 HEPES, and 2 K_2_ATP; pH 7.2.

GABA concentration response relationships were constructed by measuring currents (*I*) elicited at each GABA concentration and normalising these currents to the maximal response (*I*_*max*_). The concentration response relationship was fitted with the Hill equation:

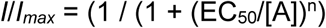

where A is GABA concentration, EC_50_ is half-maximal GABA concentration and n is the Hill slope. The macroscopic kinetics of GABA-activated currents were studied in HEK-293 cells by applying 1 mM GABA^57^. The activation rate was the time taken to ascend from 10 - 90% of *I*_*max*_ and the deactivation rate was the weighted tau of exponential fits from the point of cessation of GABA application until the baseline was attained.

I-V relationships were constructed by stepping the holding potential from −80 to 80 mV in increments of 10 mV in control and in the presence of 1 mM picrotoxin. The waveform in the presence of picrotoxin was subtracted from the basal curve to give the I-V relationship of picrotoxin-sensitive currents.

Neurons transfected at 7 *DIV* were voltage clamped using the same CsCl internal at –60 mV for recording spontaneous inhibitory postsynaptic currents (sIPSCs) and tonic currents. Neurons were superfused with the same saline solution as HEK-293 cells but supplemented with 2 mM kynurenic acid to block excitatory neurotransmission, as necessary. Membrane capacitance was measured by applying brief −10 mV hyperpolarising pulses and calculating the area under the capacity current discharge curve. Current densities were measured by dividing maximal GABA currents by the determined cell membrane capacitance.

Miniature excitatory postsynaptic currents (mEPSCs) were recorded at −70 mV in the same saline solution as HEK-293 cells but supplemented with 0.5 µM tetrodotoxin, 25 µM bicuculline and 50 µM picrotoxin using an internal solution containing (mM): 145 Cs methanesulfonate, 5 MgATP, 10 BAPTA, 0.2 Na_2_GTP, 10 HEPES, 2 QX314 and pH - 7.2.

### Imaging of dendritic spines

Dendritic spines images were collected from eGFP co-expressing live transfected neurons at 12-16 *DIV* in a saline solution containing (in mM): 140 NaCl, 4.7 KCl, 1.2 MgCl_2_, 2.52 CaCl_2_, 11 Glucose, and 5 HEPES; pH 7.4. For neurons with stereotypical pyramidal morphology, the segment of the apical dendrite closest to the soma was chosen for imaging and for neurons with non-pyramidal morphology the thickest dendrite was selected. 3D stacks of eGFP-filled dendrites were imaged with optimal z-thickness in 8-bit using a Zeiss LSM 510 microscope and a x40 water objective with an optical zoom of x2 and a 488 nm laser. Dendritic spines were analysed using Neuronstudio^58^ (Ver 0.9.92).

## References

1. Sigel, E. & Steinmann, M. E. Structure, Function, and Modulation of GABA_A_Receptors. J. Biol. Chem. 287, 40224–40231 (2012).

2. Farrant, M. & Nusser, Z. Variations on an inhibitory theme: phasic and tonic activation of GABA_A_ receptors. Nat. Rev. Neurosci. 6, 215–229 (2005).

3. Tang, X., Jaenisch, R. & Sur, M. The role of GABAergic signalling in neurodevelopmental disorders. Nat. Rev. Neurosci. 22, 290–307 (2021).

4. Möhler, H. GABA_A_ receptors in central nervous system disease: Anxiety, epilepsy, and insomnia. J. Recept. Signal Transduct. 26, 731–740 (2006).

5. Smart, T. G. & Stephenson, F. A. A half century of γ-aminobutyric acid. Brain Neurosci. Adv. 3, 239821281985824 (2019).

6. Scheffer, I. E. & Berkovic, S. F. The genetics of human epilepsy. Trends Pharmacol.Sci. 24, 428–433 (2003).

7. Wang, J. et al. Epilepsy-associated genes. Seizure 44, 11–20 (2017).

8. Audenaert, D. et al. A novel GABRG2 mutation associated with febrile seizures. Neurology 67, 687–690 (2006).

9. Hernandez, C. C. et al. Deleterious rare variants reveal risk for loss of gabaa receptor function in patients with genetic epilepsy and in the general population. PLoS One 11, e0162883 (2016).

10. Lachance-Touchette, P. et al. Novel α1 and γ2 GABA_A_ receptor subunit mutations in families with idiopathic generalized epilepsy. Eur. J. Neurosci. 34, 237–49 (2011).

11. Hannan, S. et al. Differential coassembly of α1-GABA_A_Rs associated with epileptic encephalopathy. J. Neurosci. 40, 5518–5530 (2020).

12. Hales, T. G. et al. The epilepsy mutation, γ2(R43Q) disrupts a highly conserved inter-subunit contact site, perturbing the biogenesis of GABA_A_ receptors. Mol. Cell. Neurosci. 29, 120–7 (2005).

13. Tian, M. et al. Impaired surface αβγ GABA_A_ receptor expression in familial epilepsy due to a GABRG2 frameshift mutation. Neurobiol. Dis. 50, 135–141 (2013).

14. Kang, J.-Q.J. & Macdonald, R. L. The GABA_A_ receptor γ2 subunit R43Q mutation linked to childhood absence epilepsy and febrile seizures causes retention of α1β2γ2S receptors in the endoplasmic reticulum. J. Neurosci. 24, 8672–7 (2004).

15. Sancar, F. & Czajkowski, C. A GABA_A_ receptor mutation linked to human epilepsy (γ2R43Q) impairs cell surface expression of αβγ receptors. J. Biol. Chem. 279, 47034–47039 (2004).

16. Hernandez, C. C. & Macdonald, R. L. A Structural look at GABA_A_ receptor mutations linked to epilepsy syndromes. Brain Res. 1714, 234–247 (2019).

17. Maljevic, S. et al. Spectrum of GABA_A_ receptor variants in epilepsy. Curr. Opin. Neurol. 32, 183–190 (2019).

18. Reddy, D. S. & Rogawski, M. A. Neurosteroid replacement therapy for catamenial epilepsy. Epilepsy Mech. Model. Transl. Perspect. 6, 501–513 (2010).

19. Nohria, V. & Giller, E. Ganaxolone. Neurotherapeutics 4, 102–105 (2007).

20. Hernandez, C. C. et al. GABA_A_Receptor Coupling Junction and Pore GABRB3 Mutations are Linked to Early-Onset Epileptic Encephalopathy. Sci. Rep. 7, 1–18 (2017).

21. Absalom, N. L. et al. Gain-of-function GABRB3 variants identified in vigabatrin-hypersensitive epileptic encephalopathies. Brain Commun. 2, 1–16 (2020).

22. Whiting, P. J., McKernan, R. M. & Wafford, K. A. Structure and pharmacology of vertebrate GABA_A_ receptor subtypes. Int. Rev. Neurobiol. 38, 95–138 (1995).

23. Hosie, A. M., Wilkins, M. E., da Silva, H. M. A. & Smart, T. G. Endogenous neurosteroids regulate GABA_A_ receptors through two discrete transmembrane sites. Nature 444, 486–489 (2006).

24. Hosie, A. M., Dunne, E. L., Harvey, R. J. & Smart, T. G. Zinc-mediated inhibition of GABA_A_ receptors: discrete binding sites underlie subtype specificity. Nat.Neurosci. 6, 362–369 (2003).

25. Wong, M. & Guo, D. Dendritic spine pathology in epilepsy: Cause or consequence? Neuroscience 251, 141–150 (2013).

26. Seljeset, S., Laverty, D. & Smart, T. G. Inhibitory neurosteroids and the GABA_A_ receptor. Adv. Pharmacol. 72, 165–187 (2015).

27. Jacob, T. C. et al. GABA_A_ receptor membrane trafficking regulates spine maturity. Proc. Natl. Acad. Sci. 106, 12500–12505 (2009).

28. Farnaes, L. et al. Rapid whole-genome sequencing identifies a novel GABRA1 variant associated with West syndrome. Cold Spring Harb. Mol. case Stud. 3, 1–8 (2017).

29. Kodera, H. et al. De novo GABRA1 mutations in Ohtahara and West syndromes. Epilepsia 57, 566–573 (2016).

30. Hosie, A. M., Wilkins, M. E. & Smart, T. G. Neurosteroid binding sites on GABA_A_ receptors. Pharmacol. Ther. 116, 7–19 (2007).

31. Myers, C. T. et al. De Novo Mutations in SLC1A2 and CACNA1A Are Important Causes of Epileptic Encephalopathies. Am. J. Hum. Genet. 99, 287–298 (2016).

32. Maljevic, S. et al. Novel GABRA2 variants in epileptic encephalopathy and intellectual disability with seizures. Brain 142, 1–6 (2019).

33. Bianchi, M. T. & MacDonald, R. L. Neurosteroids shift partial agonist activation of GABA_A_ receptor channels from low-to high-efficacy gating patterns. J.Neurosci. 23, 10934–10943 (2003).

34. MacKenzie, G. & Maguire, J. Neurosteroids and GABAergic signaling in health and disease. Biomol. Concepts 4, 29–42 (2013).

35. Jaslove, S. W. The integrative properties of spiny distal dendrites. Neuroscience 47, 495–519 (1992).

36. Nimchinsky, E. A., Sabatini, B. L. & Svoboda, K. Structure and Function of Dendritic Spines. Annu. Rev. Physiol. 64, 313–353 (2002).

37. Kasai, H., Fukuda, M., Watanabe, S., Hayashi-Takagi, A. & Noguchi, J. Structural dynamics of dendritic spines in memory and cognition. Trends Neurosci. 33, 121–129 (2010).

38. Fernandez, F. & Garner, C. C. Over-inhibition: a model for developmental intellectual disability. Trends Neurosci. 30, 497–503 (2007).

39. Fernandez, F. et al. Pharmacotherapy for cognitive impairment in a mouse model of Down syndrome. Nat Neurosci 10, 411–413 (2007).

40. Belichenko, P. V et al. Synaptic structural abnormalities in the Ts65Dn mouse model of Down Syndrome. J.Comp Neurol. 480, 281–298 (2004).

41. Iwai, Y., Fagiolini, M., Obata, K. & Hensch, T. K. Rapid critical period induction by tonic inhibition in visual cortex. J. Neurosci. 23, 6695–6702 (2003).

42. Majewska, A. & Sur, M. Motility of dendritic spines in visual cortex in vivo: Changes during the critical period and effects of visual deprivation. Proc. Natl. Acad. Sci. U. S. A. 100, 16024–16029 (2003).

43. Mataga, N., Mizuguchi, Y. & Hensch, T. K. Experience-dependent pruning of dendritic spines in visual cortex by tissue plasminogen activator. Neuron 44, 1031–1041 (2004).

44. Hensch, T. K. Critical period plasticity in local cortical circuits. Nat. Rev. Neurosci. 6, 877–888 (2005).

45. Hensch, T. K. & Fagiolini, M. Excitatory-inhibitory balance and critical period plasticity in developing visual cortex. Prog. Brain Res. 147, 115–124 (2005).

46. Hayama, T. et al. GABA promotes the competitive selection of dendritic spines by controlling local Ca^2+^ signaling. Nat Neurosci 16, 1409–1416 (2013).

47. Wong, M. Modulation of dendritic spines in epilepsy: Cellular mechanisms and functional implications. Epilepsy Behav. 7, 569–577 (2005).

48. Swann, J. W., Al-Noori, S., Jiang, M. & Lee, C. L. Spine loss and other dendritic abnormalities in epilepsy. Hippocampus 10, 617–625 (2000).

49. Hasbani, M. J., Hyrc, K. L., Faddis, B. T., Romano, C. & Goldberg, M. P. Distinct roles for sodium, chloride, and calcium in excitotoxic dendritic injury and recovery. Exp. Neurol. 154, 241–258 (1998).

50. Al-Noori, S. & Swann, J. W. A role for sodium and chloride in kainic acid-induced beading of inhibitory interneuron dendrites. Neuroscience 101, 337–348 (2000).

51. Adrian, M. et al. Barriers in the brain: Resolving dendritic spine morphology and compartmentalization. Front. Neuroanat. 8, 1–12 (2014).

52. Gulledge, A. T., Carnevale, N. T. & Stuart, G. J. Electrical advantages of dendritic spines. PLoS One 7, e36007 (2012).

53. Laverty, D. et al. Cryo-EM structure of the human α1β3γ2 GABA_A_ receptor in a lipid bilayer. Nature 565, 516–520 (2019).

54. Hannan, S., Wilkins, M. E., Thomas, P. & Smart, T. G. Tracking cell surface mobility of GPCRs using alpha-bungarotoxin-linked fluorophores. Methods Enzym. 521, 109–129 (2013).

55. Hannan, S. et al. GABA_A_R isoform and subunit structural motifs determine synaptic and extrasynaptic receptor localisation. Neuropharmacology 169, 107540 (2019).

56. Hannan, S., Wilkins, M. E. & Smart, T. G. Sushi domains confer distinct trafficking profiles on GABA_B_ receptors. Proc.Natl.Acad.Sci.U.S.A 109, 12171–12176 (2012).

57. Thomas, P. & Smart, T. G. Use of electrophysiological methods in the study of recombinant and native neuronal ligand-gated ion channels. Curr. Protoc. Pharmacol. (2012) doi:10.1002/0471141755.ph1104s59.

58. Rodriguez, A., Ehlenberger, D. B., Dickstein, D. L., Hof, P. R. & Wearne, S. L. Automated three-dimensional detection and shape classification of dendritic spines from fluorescence microscopy images. PLoS One 3, e(1997) (2008).

