## Supplementary Data for "Inhibitory neurosteroid reverses the dendritic spine disorder caused by gain-of-function GABA_A_R epilepsy variants"

Supplementary Fig. 1

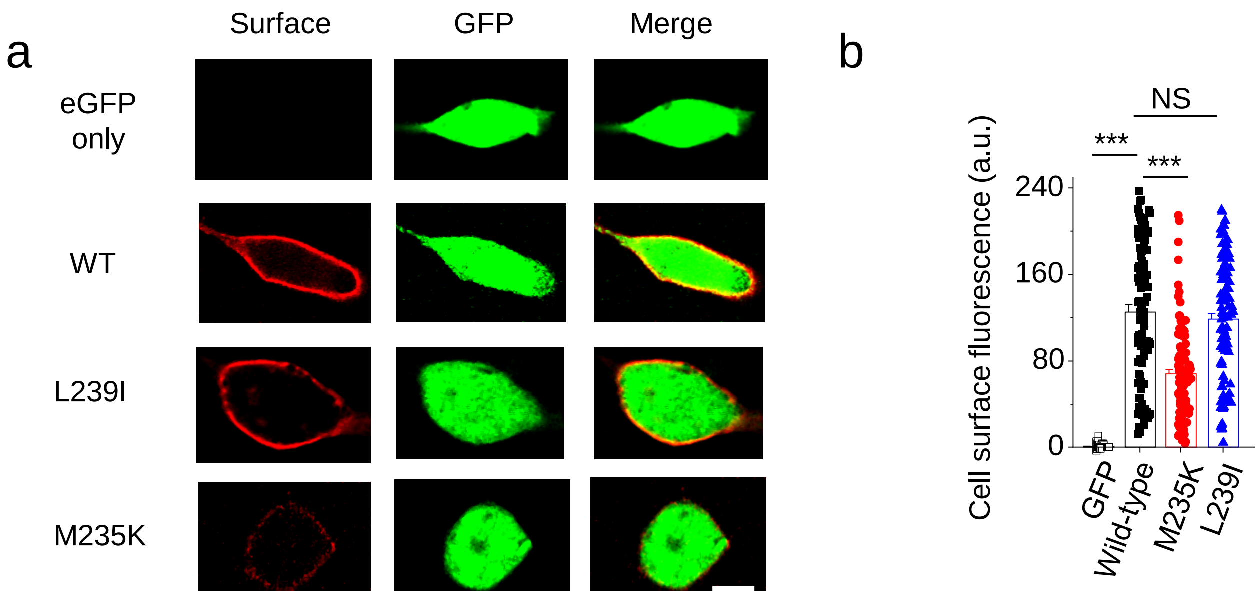

#### Supplementary Fig. 1. Cell surface expression of $GABA_A$Rs in HEK-293 cells

**a** Confocal images of cell surface α1 wild-type (WT) and α1-variant $GABA_A$Rs expressed in HEK-293 cells. Cells expressing eGFP with or without wild-type or variant $α1^{myc}$, β2 and γ2L were immunolabelled with anti-myc antibodies prior to imaging. **b** Mean cell surface fluorescence intensities (arbitrary units, a.u.) for WT and α1-variant $GABA_A$Rs. n = 97-105 cells, NS – not significant, ***$P<0.001$, One-way ANOVA. $F_{(3, 394)} = 140.81$, $p<0.0001$. Calibration bars = 5 μm. Bars are mean +/- SEM, including individual data points.

### Supplementary Fig. 2

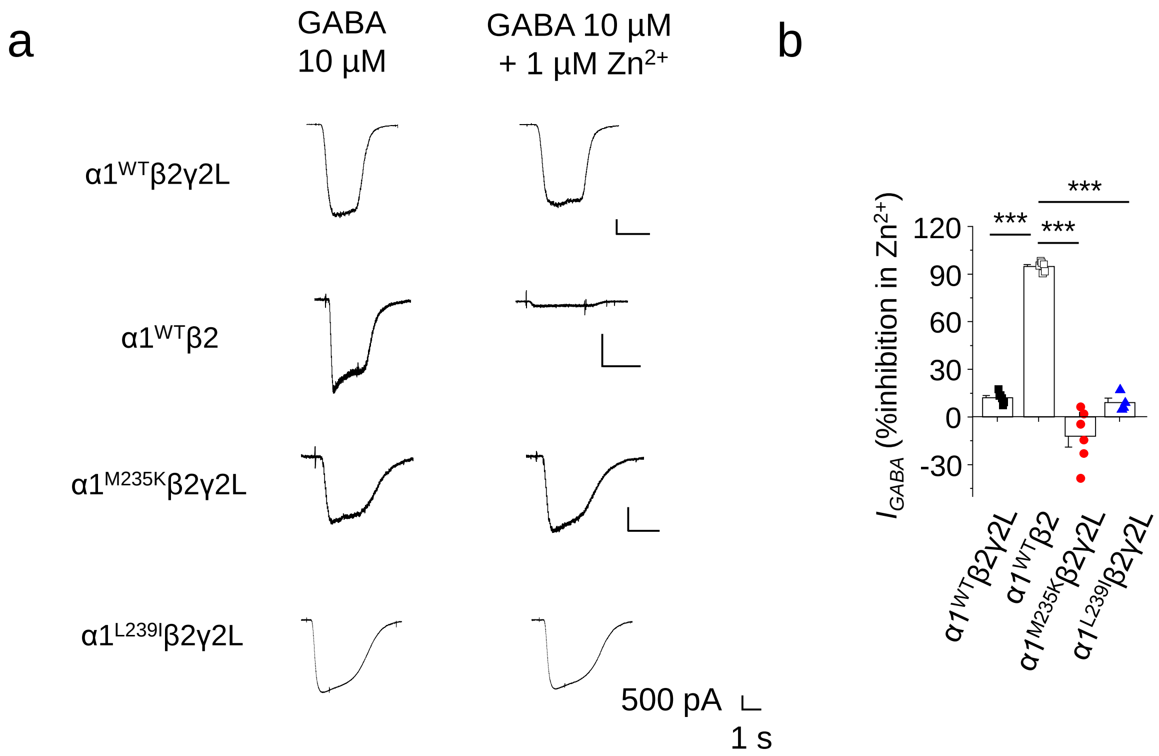

#### Supplementary Fig. 2. $Zn^{2+}$ inhibition at $GABA_A$R variants

**a** Whole-cell 10 µM GABA-activated currents recorded in HEK-293 cells in control or in the presence of 1 µM $Zn^{2+}$ for HEK cells expressing wild-type or variant α1, β2, γ2L and eGFP, or wild-type α1, β2 and GFP. 1 µM $Zn^{2+}$ was pre-applied for 5 - 15 s. **b** % Inhibition of GABA-activated currents by 1 µM $Zn^{2+}$. $F_{(3, 18)} = 144.8$, $p<0.0001$, n = 4 – 5 cells. ***$P<0.001$, One-way ANOVA.

### Supplementary Fig. 3

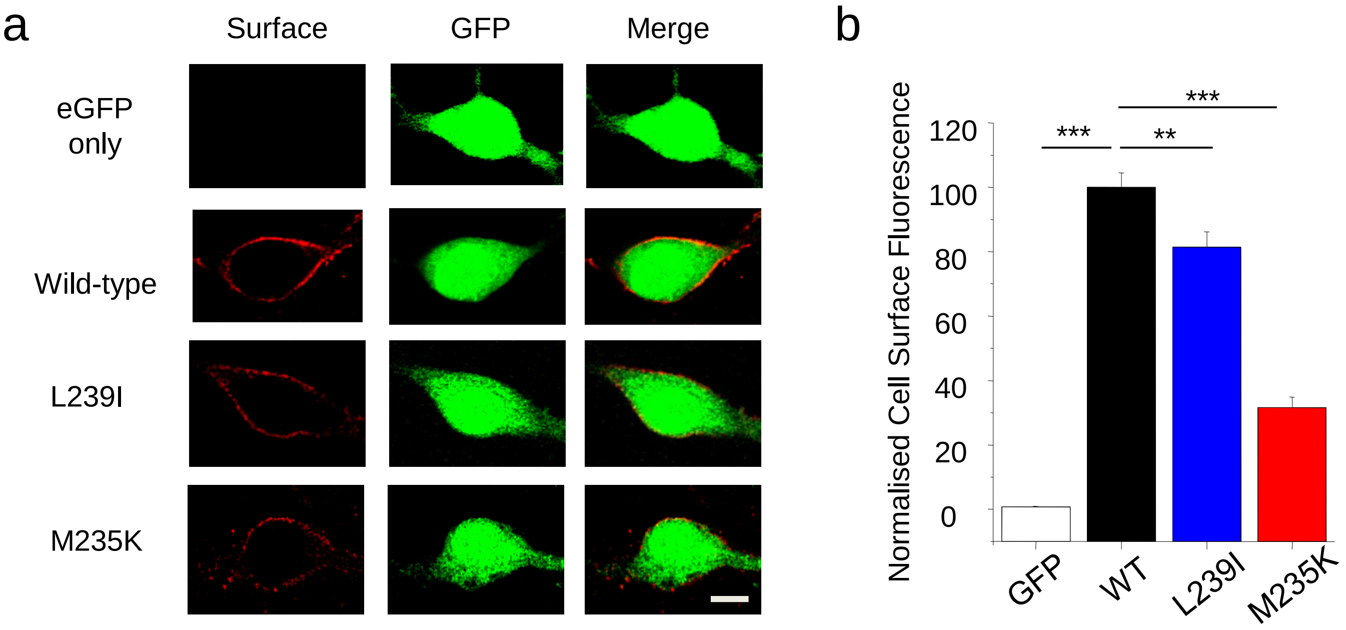

#### Supplementary Fig. 3. Cell surface expression of $GABA_A$Rs in neurons

**a** Confocal images of cell surface expression for α1 wild-type and α1-variant $GABA_A$Rs for hippocampal neurons in culture. Neurons expressing α1 wild-type or variant $α1^{myc}$ and eGFP were immunolabelled with anti-myc antibodies prior to imaging. **b** Normalised (%) mean cell surface fluorescence intensities for α1 wild-type (WT) and α1-variant $GABA_A$Rs. Data were normalised to wild-type expression levels. $F_{(3, 167)}$ = 136.5, p<0.0001, n = 39 - 46 neurons, **P<0.01, ***P<0.001, One-way ANOVA. Calibration bar = 5 μm.

Supplementary Fig. 4

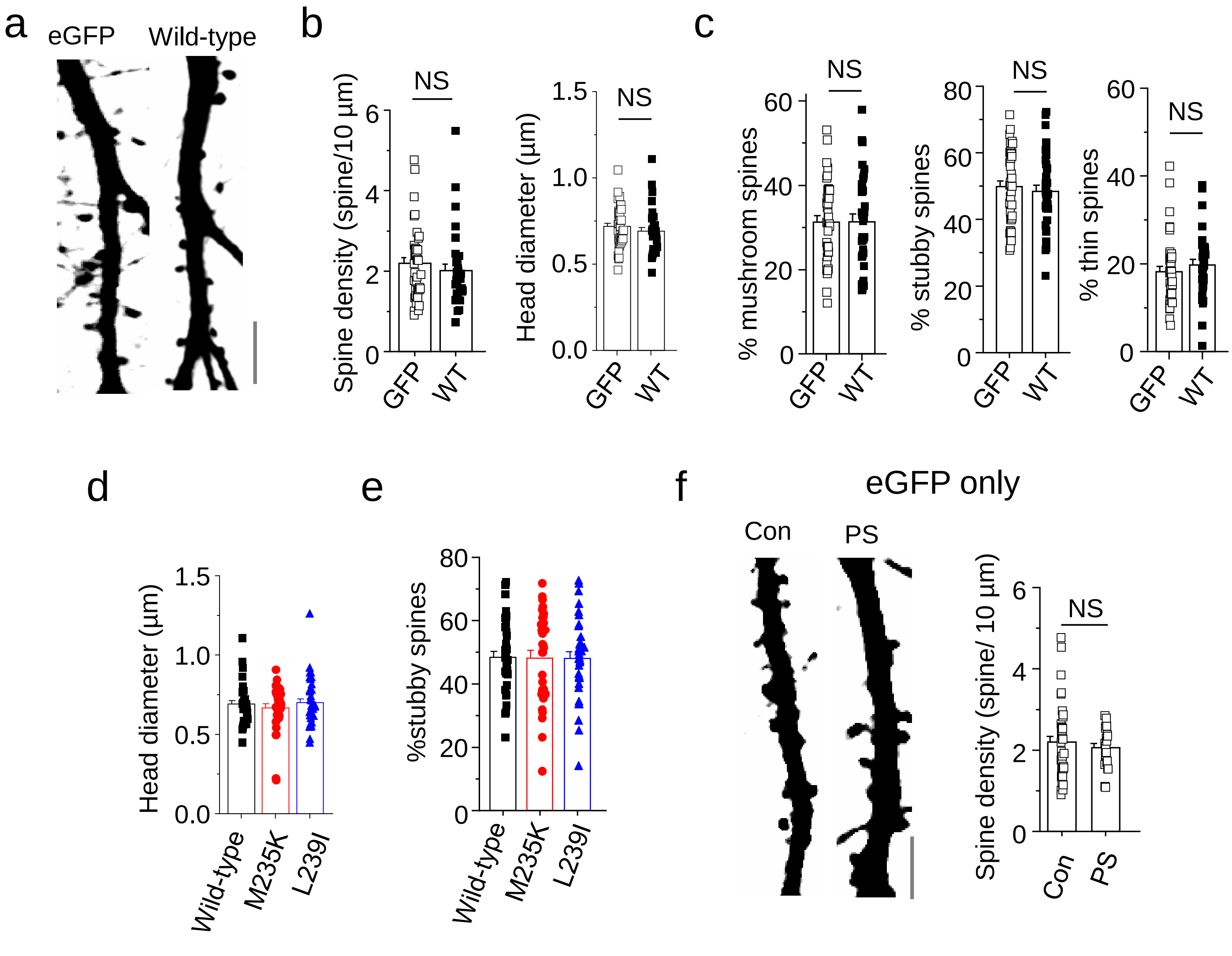

#### Supplementary Fig. 4. Dendritic spine properties after α1-$GABA_A$R expression and effect of pregnenolone sulphate in GFP-expressing neurons

**a** Confocal images of dendrites from hippocampal neurons expressing eGFP only or eGFP with wild-type α1-$GABA_A$Rs. **b** Mean spine density (p=0.2481) and head diameter (p=0.3413) of wild-type α1-$GABA_A$R (WT) expressing neurons or eGFP (GFP) only controls. **c** Proportions (%) of mushroom-shaped (p=0.966), stubby (p=0.6568) and thin (p=0.4366) spines of wild-type α1-$GABA_A$R (WT) expressing neurons or eGFP (GFP) only controls. **d** Bargraph of spine head diameter for wild-type and α1-variant $GABA_A$Rs in cultured hippocampal neurons. $F_{(2,104)}$ = 0.4776, p=0.6217. **e** Bargraph of mean proportion (%) of stubby spines for wild-type and α1-variant $GABA_A$R expressing hippocampal neurons in culture. $F_{(2,101)}$ = 0.019, p=0.9816. **f** Confocal images of dendrites from hippocampal neurons expressing eGFP only in control (con) and after neurons were treated with 5 μM pregnenolone sulphate (PS) for 48 hr at 37°C (p = 0.9005). n = 34 - 40 neurons. NS- not significant, one-way ANOVA, two-tailed unpaired t-test. Calibration bar = 5 μm.

### Supplementary Fig. 5

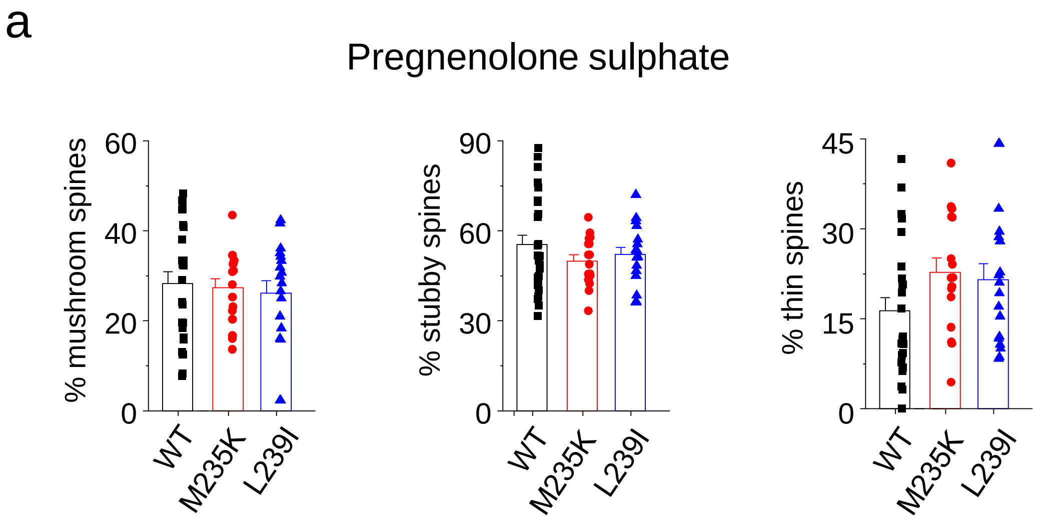


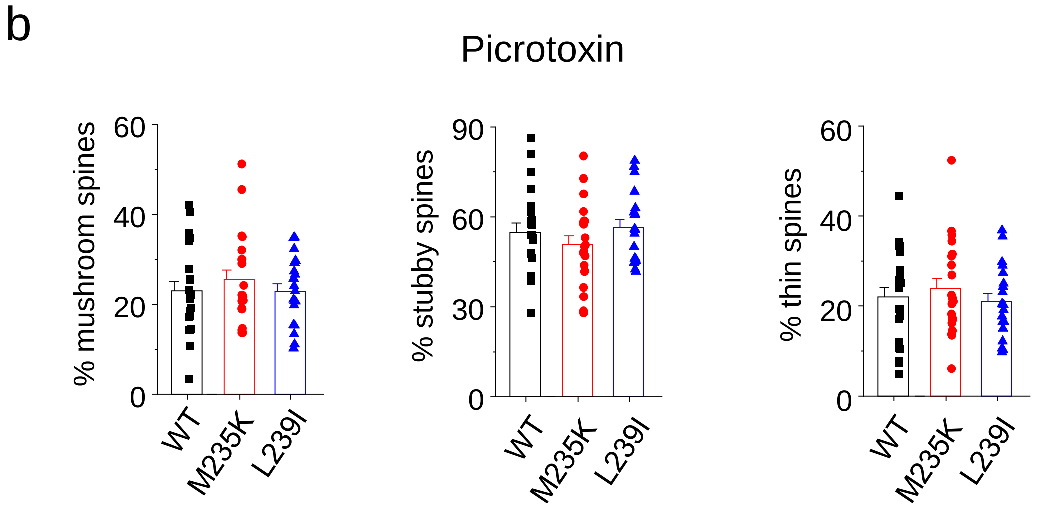

#### Supplementary Fig. 5. Dendritic spine morphology in pregnenolone sulphate and picrotoxin-treated neurons

**a** Proportion (%) of mushroom-shaped, stubby and thin spines of hippocampal neurons expressing α1 wild-type, $\alpha1^{M235K}$ or $\alpha1^{L239I}$ in pregnenolone sulphate. Neurons were treated with 5 μM pregnenolone sulphate for 48 hr at 37°C before imaging. $F_{(2, 57)}$ = 0.189, p = 0.8281 for mushroom spines; $F_{(2, 57)}$ = 0.984, p = 0.3801 for stubby spines; $F_{(2, 56)}$ = 2.05, p = 0.1385 for thin spines. **b** Proportion of mushroom-shaped, stubby and thin spines of hippocampal neurons expressing α1 wild-type, $\alpha1^{M235K}$ or $\alpha1^{L239I}$ with eGFP in picrotoxin. Neurons were treated with 50 μM picrotoxin for 48 hr at 37°C prior to imaging. $F_{(2, 59)}$ = 0.504, p = 0.6066 for mushroom spines; $F_{(2, 59)}$ = 0.9838, p = 0.3799 for stubby spines; $F_{(2, 59)}$ = 0.4448, p = 0.6431 for thin spines. n = 16 - 40 neurons. One-way ANOVA.
